# Brain state stability during working memory is explained by network control theory, modulated by dopamine D1/D2 receptor function, and diminished in schizophrenia

**DOI:** 10.1101/679670

**Authors:** Urs Braun, Anais Harneit, Giulio Pergola, Tommaso Menara, Axel Schaefer, Richard F. Betzel, Zhenxiang Zang, Janina I. Schweiger, Kristina Schwarz, Junfang Chen, Giuseppe Blasi, Alessandro Bertolino, Daniel Durstewitz, Fabio Pasqualetti, Emanuel Schwarz, Andreas Meyer-Lindenberg, Danielle S. Bassett, Heike Tost

## Abstract

Dynamical brain state transitions are critical for flexible working memory but the network mechanisms are incompletely understood. Here, we show that working memory entails brain-wide switching between activity states. The stability of states relates to dopamine D1 receptor gene expression while state transitions are influenced by D2 receptor expression and pharmacological modulation. Schizophrenia patients show altered network control properties, including a more diverse energy landscape and decreased stability of working memory representations.

---

Working memory is an essential part of executive cognition depending on prefrontal neurons functionally modulated through dopamine D1 and D2 receptor activation (1-3). The dual-state theory of prefrontal dopamine function links the differential activation of dopamine receptors to two discrete dynamical regimes: a D1-dominated state with a high energy barrier favoring robust maintenance of cognitive representations and a D2-dominated state with a flattened energy landscape enabling flexible switching between states (4). Recent accounts extend the idea of dopamine’s impact on working memory from a local prefrontal to a brain-wide network perspective (5, 6), but the underlying neural dynamics and brain-wide interactions have remained unclear.

Network control theory (NCT) can be used to model brain network dynamics as a function of interconnecting white matter tracts and regional control energy (7). Based on the connectome, NCT can be used to examine the landscape of brain activity states: that is, which states within a dynamic scheme would the system have difficulty accessing, and more importantly, which regions need to be influenced (and to what extent) to make those states accessible (8). Specifically, to quantify accessibility, we approximate brain dynamics locally by a simple linear dynamical system, 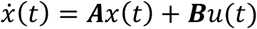, where *x (t)* is the brain state inferred from functional magnetic resonance imaging (fMRI), ***A*** is a structural connectome inferred from DTI data, *u* is the control input, and ***B*** is a matrix describing which regions enact control. To investigate, based on this conception, how the brain transitions between different cognitive states, we defined states as individual brain activity patterns related to a working memory condition (2-back) and to an attention control condition requiring motor response (0-back) in a sample of 178 healthy individuals undergoing fMRI (***Fig. S1;*** Online Methods). Further, we obtained individual structural connectomes from white matter by DTI fiber tracking, and computed the optimal control energy necessary to drive the dynamical system from the 0-back activity pattern to the 2-back pattern, or *vice versa* (***Fig. S2***).

We defined the stability of both brain states as the inverse energy necessary to revisit that state, where the energy, loosely, is defined as the average size of the control signals *u(t*) needed to instantiate a specific trajectory in the dynamical system as defined above (see Eq. 3 & 5 in Online Methods). As expected, the cognitively more demanding 2-back state was less stable (i.e., required higher energy for maintenance) than the control state (***Fig. 1a***; repeated measures ANOVA: main effect of 0-vs. 2-back stability: F(1,173) = 66.80, p < 0.001, see Online Methods for details on all analyses). Further, the stability of the 2-back state was significantly associated with working memory accuracy (***Fig. 1b***; *b* = 0.274, p = 0.006), suggesting that more stable 2-back network representations support higher working memory performance. We next investigated how the brain flexibly changes its activity pattern between states. Transitioning into the cognitively more demanding 2-back state required more control energy than the opposite transition (***Fig. 1c***; repeated measures ANOVA: F(1,174) = 27.98, p = 0.001). Other analyses suggested that prefrontal and parietal cortices steer both types of transitions, while default mode areas are preferentially important for the switch to the more cognitively demanding state (***Fig. 1d***; Online Methods). These results are in line with the assumed role of frontal-parietal circuits in steering brain dynamics (9) and shifting brain connectivity patterns (10); they also emphasize the importance of the coordinated behavior of brain systems commonly displaying deactivations during demanding cognitive tasks (11).

**Figure 1:**
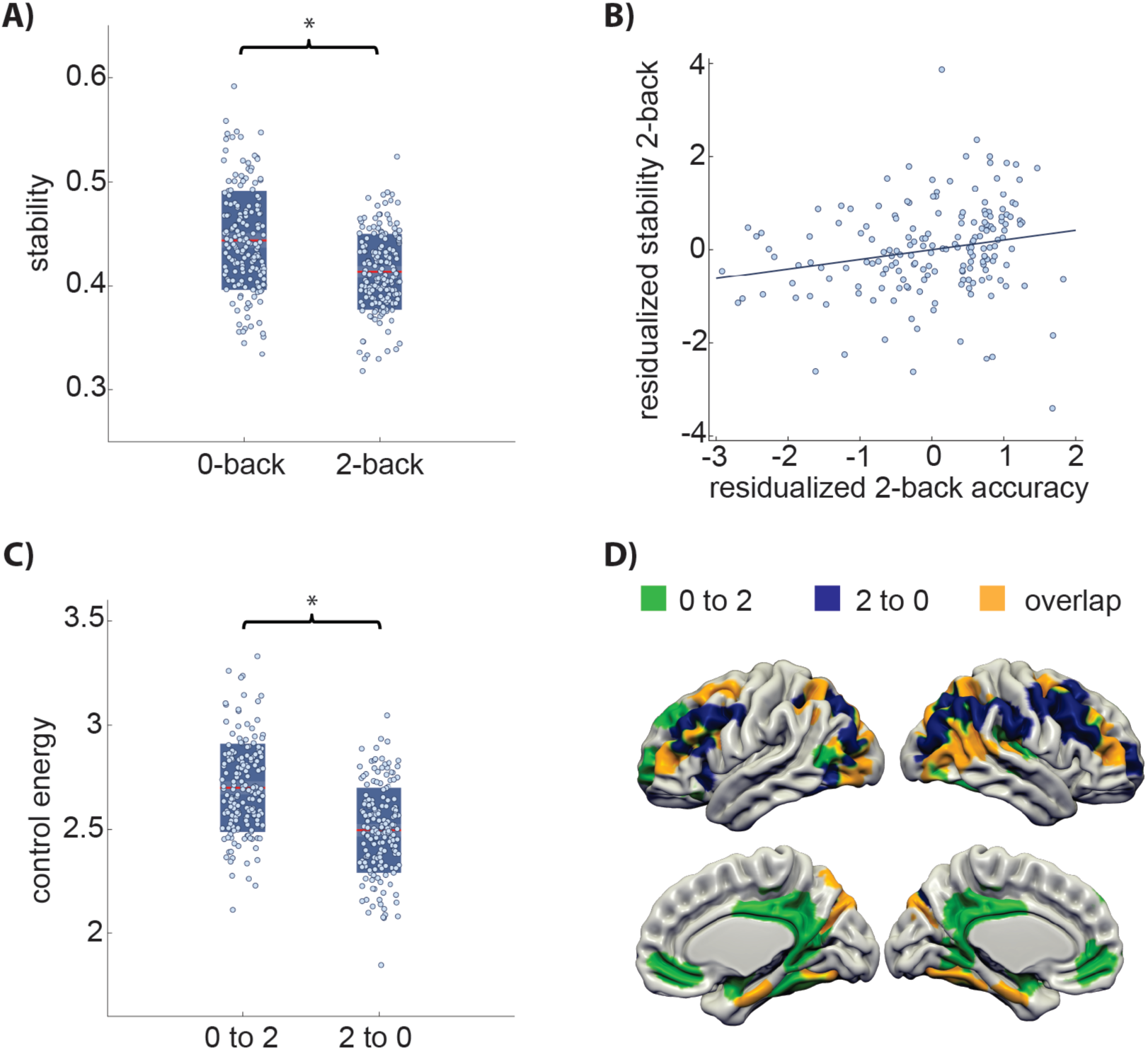
Controllability and stability of brain dynamics during working memory. A) The stability of the 2-back state reflecting working memory activity is lower than that of the 0-back state reflecting motor and basic attention control activity (F(1,173) = 66.80, p < 0.001). Red lines indicate mean values and boxes indicate one standard deviation of the mean. B) Associations of 2-back stability with working memory performance (accuracy: *b* = 0.274, p = 0.006; covarying for age, sex, and mean activity). C) Steering brain dynamics from the control condition to the working memory condition requires more control energy than *vice versa* (F(1,174) = 27.98, p < 0.001). D) Unique and common sets of brain regions contribute most to the transition from 0-back to 2-back and the transition from 2-back to 0-back transitions, respectively. For illustrative purposes, we projected the computed control impact of each brain region (Online Methods) for the respective transitions on a 3D structural template, displaying the 20% highest for each transition.

Following from the dual-state theory of network function, the stability of task-related brain states should be related to prefrontal D1 receptor status. To estimate individual prefrontal D1 receptor expression, we utilized methods relating prefrontal cortex D1 and D2 receptor expression to genetic variation in their co-expression partner (Online Methods), thereby enabling us to predict individual dopamine receptor expression levels from genotype data across the whole genome (12, 13). We found that D1 (but not the D2) expression-related gene score predicted stability of both states (***Fig. 2a***; 0-back: *b* = 0.184, p = 0.034; 2-back: *b* = 0.242, p = 0.007, Online Methods), in line with the assumed role of D1-related signaling in maintaining stable activity patterns during task performance (4, 14).

**Figure 2:**
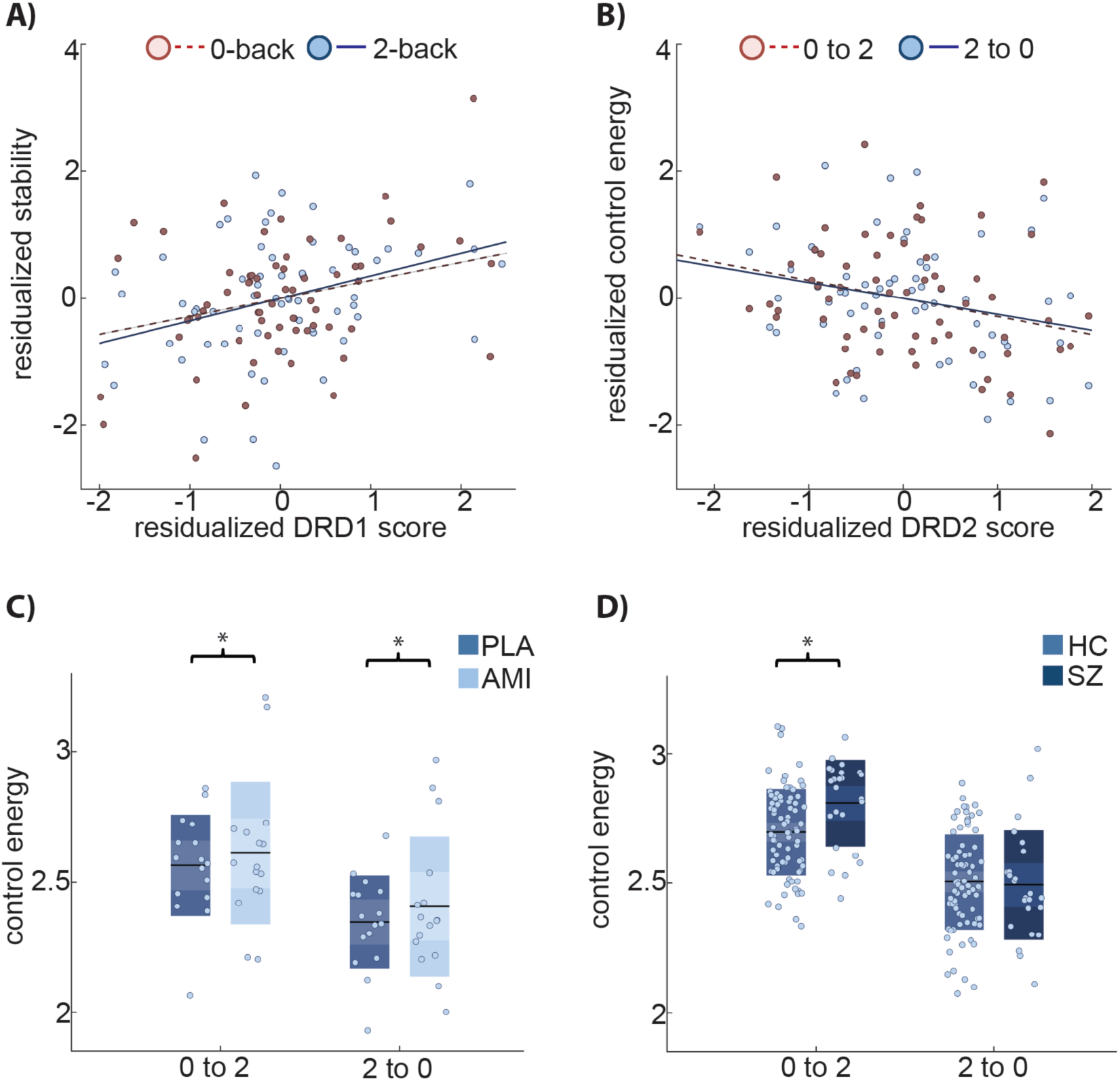
Dopamine receptor expression and pharmacological modulation impact whole brain dynamics. A) Genetic scores predicting DRD1 expression in prefrontal regions positively predict stability of both brain states (0-back: *b* = 0.184, p = 0.034; 2-back: *b* = 0.242 p = 0.007; age, sex, mean brain state activity, first 5 genetic PCA components as covariates of non-interest). B) Genetic scores predicting DRD2 expression in prefrontal regions negatively predict control energy for both brain state transitions (0-back to 2-back: *b* = −0.076, p = 0.037; and trend wise for 2-back to 0-back: *b* = −0.134, p = 0.068; age, sex, mean brain activity difference, first 5 genetic PCA components, stability of 0-back and 2-back as covariates of non-interest). C) Amisulpride increases control energy for transitions in comparison to placebo (main effect of drug: F(1,10) = 7.27, p = 0.022; interaction drug by condition: F(1,10) = 0.42, p = 0.665, activity difference, drug order, and sex as covariates of non-interest). Black lines indicate mean values and boxes indicate one standard deviation of the mean. D) Schizophrenia patients need more control energy when transitioning into the working memory condition than matched healthy controls (F(1,98) = 5.238, p = 0.024, age, sex, tSNR and mean activity as covariates of non-interest), but not *vice versa*.

Independent of stability, switching between different activity representations should relate to dopamine D2 receptor function. Indeed, when controlling for stability as a nuisance covariate in the regression model, the control energy of both state transitions could be predicted by the D2 (but not the D1) receptor expression gene score (***Fig. 2b***; 0- to 2-back: *b* = −0.076, p = 0.037; and trending for 2- to 0-back: *b* = −0.134, p = 0.068, Online Methods). This finding is particularly interesting, as it suggests that the function of D1 and D2 receptors are differentially, but cooperatively, involved in steering brain dynamics between different activity patterns, in line with previous research on D1 and D2 functioning in prefrontal circuits (4, 15).

Our results thus far support the notion that the brain is a dynamical system in which the stability of a state is substantially defined by cognitive effort and modulated by D1 receptor expression, while transitions between states depend primarily on D2 receptor expression. If true, such a system should be sensitive to dopaminergic manipulation, and interference with D2-related signaling should reduce the brain’s ability to control its optimal trajectories, i.e. increase the control energy needed when switching between states. To test these hypotheses, we investigated an independent sample of healthy controls (n=16, ***Table S2***) receiving 400 mg Amisulpride, a selective D2 receptor antagonist, in a randomized, placebo-controlled, double-blind pharmacological fMRI study. As expected, we observed that greater control energy was needed for transitions under D2 receptor blockade (***Fig. 2c***; repeated measures ANOVA with drug and transition as within-subject factors; main effect of drug: F(1,10) = 7.27, p = 0.022; drug-by-condition interaction: F(1,10) = 0.42, p = 0.665). We observed no effect on the stability of states; that is, the inverse control energy required to stabilize a current state (main effect of drug: F(1,8) = 0.715, p = 0.422, ***Table S3***).

Dopamine dysfunction, working memory deficits, and alterations in brain network organization are hallmarks of schizophrenia (16-19). We therefore tested for differences in the state stability and in the ability to control state transitions between schizophrenia patients and a healthy control sample balanced for age, sex, performance, head motion, and premorbid IQ (see ***Table S1***). Stability in schizophrenia patients was reduced for the cognitively demanding working memory state (F(1,98) = 6.43, p = 0.013), but not for the control condition (F(1,98) = 0.052, p = 0.840, ***Table S3***). Control energy needed for the 0- to 2-back transition was significantly higher in schizophrenia (***Fig. 2d***; F(1,98) = 5.238, p = 0.024), while the opposite transition showed no significant group difference (ANOVA: F(1,98) = 0.620, p = 0.433, ***Table S3***), in line with clinical observations that D2 blockade does not ameliorate cognitive symptoms in schizophrenia (20). These results suggest that the brain energy landscape is more diverse in schizophrenia, making the system more difficult to steer appropriately. To further strengthen this notion, we estimated the variability in suboptimal (higher energy) trajectories connecting different of cognitive states (Online Methods). We expected that in a diversified energy landscape, the variation of trajectories around the minimum-energy trajectory should be larger, implying that small perturbations may have a more substantial impact. In line with our hypothesis, we found that the variability in such perturbed trajectories was indeed increased in schizophrenia (rm-ANOVA: main effect of group: F(1,98) = 4.789, p = 0.031, Online Methods).

Several aspects of our work require special consideration. Firstly, to relate brain dynamics to cognitive function, we focus on discrete brain states where each state is summarized by a single brain activation patterns rather than linear combination of multiple brain activity patterns. Secondly, although we could demonstrate a link between brain dynamics, measured by means of control energy, and predicted prefrontal dopamine receptor expression, the link is indirect and requires confirmation by direct measurements. Thirdly, we cannot exclude the possibility that disorder severity, duration, symptoms or medication may have influenced network dynamics in schizophrenia patients, although our supplemental analyses do not support this conclusion (Online Methods). Finally, while the sample sizes of our pharmacological and patient study are rather small, we were able to show comparable effects of dopaminergic manipulation on control properties using a second (Online Methods), further supporting the validity of the underlying rationale.

In summary, our data demonstrate the utility of network control theory for the non-invasive investigation of the mechanistic underpinnings of (altered) brain states and their transitions during cognition. Our data suggest that engagement of working memory involves brain-wide switching between activity states and that the steering of these network dynamics is differentially, but cooperatively, influenced by dopamine D1 and D2 receptor function. Moreover, we show that schizophrenia patients show reduced controllability and stability of working memory network dynamics, consistent with the idea of an altered functional architecture and energy landscape of cognitive brain networks.

## Acknowledgements

The authors thank all individuals who have supported our work by participating in our studies. There was no involvement by the funding bodies at any stage of the study. We thank Oliver Grimm, Leila Haddad, Michael Schneider, Natalie Hess, Sarah Plier and Petya Vicheva for valuable research assistance. The authors thank Jason Kim and Lorenzo Caciagli for valuable feedback on the manuscript.

U.B. acknowledges grant support by the German Research Foundation (DFG, grant BR 5951/1-1). H.T. acknowledges grant support by the German Research Foundation (DFG, Collaborative Research Center SFB 1158 subproject B04, Collaborative Research Center TRR 265 subproject A04, GRK 2350 project B2, grant TO 539/3-1) and German Federal Ministry of Education and Research (BMBF, grants 01EF1803A project WP3, 01GQ1102). AML acknowledges grant support by the German Research Foundation (DFG, Collaborative Research Center SFB 1158 subproject B09, Collaborative Research Center TRR 265 subproject S02, grant ME 1591/4-1) and German Federal Ministry of Education and Research (BMBF, grants 01EF1803A, 01ZX1314G, 01GQ1003B), European Union’s Seventh Framework Programme (FP7, grants 602450, 602805, 115300 and HEALTH-F2-2010-241909, Innovative Medicines Initiative Joint Undertaking (IMI, grant 115008) and Ministry of Science, Research and the Arts of the State of Baden-Wuerttemberg, Germany (MWK, grant 42-04HV.MED(16)/16/1). DSB and RBF would like to acknowledge support from the John D. and Catherine T. MacArthur Foundation, the Alfred P. Sloan Foundation, the Army Research Laboratory and the Army Research Office through contract numbers W911NF-10-2-0022 and W911NF-14-1-0679, the National Institute of Health (2-R01-DC-009209-11, 1R01HD086888-01, R01-MH107235, R01-MH107703, and R21-M MH-106799), the Office of Naval Research, and the National Science Foundation (BCS-1441502, CAREER PHY-1554488, and BCS-1631550). E.S. gratefully acknowledges grant support by the Deutsche Forschungsgemeinschaft, DFG (SCHW 1768/1-1). X.L.Z. is a Ph.D. scholarship awardee of the Chinese Scholarship Council. DD acknowledges grant support by the German Research Foundation (DFG, Du 354/10-1). G.P. has received funding from the European Union’s Horizon 2020 research and innovation program under the Marie Skłodowska-Curie No. 798181: “IdentiFication of brain deveLopmental gene co-expression netwOrks to Understand RIsk for SchizopHrenia” (FLOURISH).

The content of this paper is solely the responsibility of the authors and does not necessarily represent the official views of any of the funding agencies

